# Three-Dimensional Photoacoustic Tomography with Ultrasound Localization Priors

**DOI:** 10.64898/2026.05.04.722751

**Authors:** Haoming Huo, Yirui Xu, Rui Yao, Matt Lowerison, Pengfei Song, Junjie Yao

**Affiliations:** Department of Biomedical Engineering, Duke University, Durham, NC, 27708, USA; Department of Neurology, Duke University School of Medicine, Durham, NC, 27710, USA

**Keywords:** Photoacoustic tomography, ultrasound localization microscopy, model-based reconstruction, spatial priors, super-resolution vascular imaging, functional imaging

## Abstract

Three-dimensional photoacoustic tomography (3D-PAT) enables noninvasive structural and functional imaging with optical absorption contrast and ultrasonic detection depth. However, its spatial resolution is limited by acoustic diffraction, and incomplete detection geometry can substantially degrade image fidelity and quantitative accuracy. Here, we present a ULM-guided model-based reconstruction framework, termed 3D-PAULM^prior^ that incorporates sub-diffraction vascular priors from concurrent ultrasound localization microscopy (ULM) into 3D photoacoustic reconstruction. The method uses weighted regional Laplacian regularization to integrate high-resolution vascular information into the inverse problem, thereby enhancing vascular sharpness, suppressing limited-view artifacts, and improving blood oxygen saturation estimation. We validated 3D-PAULM^prior^ using numerical simulations, tissue-mimicking phantoms, and *in vivo* mouse brain imaging. Compared with conventional reconstruction, 3D- PAULM^prior^ improved spatial resolution by over 50%, increased contrast-to-noise ratio by 261.2%, and enhanced structural similarity index by 24.6%. *In vivo*, 3D-PAULM^prior^ recovered vascular structures that were poorly resolved or missing in conventional reconstructions and produced more spatially confined sO_2_ maps. These results establish 3D-PAULM^prior^ as a robust multimodal reconstruction strategy for high-resolution structural and functional photoacoustic imaging.

## Introduction

Photoacoustic tomography (PAT) is a hybrid imaging modality that combines optical absorption contrast with ultrasonic detection, enabling high-resolution imaging at depths beyond the optical diffusion limit [1]. This unique capability has enabled a broad range of preclinical and clinical applications, including cancer diagnosis [2, 3], functional brain imaging [4-7], and hemodynamic mapping [8, 9]. Among its functional capabilities, quantification of blood oxygen saturation (sO_2_) provides important insight into tissue metabolism and pathophysiology [10].

Despite these advantages, accurate PAT reconstruction remains challenging. Image quality is often degraded by wavelength-dependent optical attenuation and incomplete acoustic detection geometry [11]. Conventional reconstruction algorithms such as delay-and-sum (DAS) are computationally efficient but prone to artifacts and resolution loss [12]. In contrast, model-based reconstruction explicitly accounts for acoustic wave propagation and improves quantitative accuracy, albeit at increased computational cost. Importantly, model-based reconstruction can incorporate prior information about the spatial distribution of absorbers, offering a practical route to mitigate the ill-posed nature of the inverse problem [13-15]. Nevertheless, in three-dimensional (3D) imaging, the reconstruction problem remains highly ill-conditioned, frequently leading to streak artifacts, reduced contrast, and loss of fine structural detail [11, 15-17].

To address these limitations, a range of regularization strategies has been proposed. Techniques such as total variation [18] and wavelet sparsity constraints [19, 20] stabilize the inversion and suppress noise, while methods such as truncated generalized singular value decomposition target limited-view artifacts [21]. However, these generic priors rely heavily on the statistical image features, but do not exploit object-specific anatomical information. To overcome this limitation, Yang et al. introduced a regional Laplacian regularization framework that integrates co-registered ultrasound (US) structural information, improving reconstruction fidelity [22]. Similar strategies have been adopted in multimodal imaging, including PET/MRI [23], FMT/CT [24], and DOT/MRI [25], where high-resolution structural priors guide reconstruction in lower-resolution modalities.

Ultrasound localization microscopy (ULM) offers a compelling source of such priors. By localizing and tracking individual microbubbles flowing across ultrafast ultrasound acquisitions, ULM achieves sub-diffraction vascular imaging with up to an order-of-magnitude improvement in spatial resolution [26-29]. Recent efforts have combined PA and ULM imaging within a single imaging platform [30]; however, these approaches have yet used the super-resolved vascular information to directly enhance PA image reconstruction.

In this work, we introduce a 3D model-based reconstruction framework that integrates ULM-derived structural priors into PAT, termed 3D-PAULM^prior^. Using a hemispherical ultrasound array, PA and ULM data are acquired in a co-registered manner. The super-resolved vascular map from ULM is incorporated into the reconstruction via a weighted regional Laplacian regularization, which accounts for the resolution mismatch and enforces spatially adaptive constraints. This formulation selectively enhances vascular features while suppressing background artifacts.

We evaluate the proposed method through simulations, phantom experiments, and *in-vivo* mouse brain imaging, and benchmark its performance against 3D-DAS and Tikhonov-regularized model-based reconstruction (3D-MBT). Multispectral acquisitions at 730, 800, and 870 nm are used to assess sO_2_ quantification. The results demonstrate consistent improvements in both structural fidelity and functional accuracy.

## Materials and Methods

### A. Model-Based Photoacoustic Reconstruction

In PAT, the acoustic pressure field is governed by the photoacoustic wave equation [31], where the pressure distribution arises from optical energy deposition following pulsed laser excitation. Under the assumption of acoustically homogeneous medium, the measured signals can be modeled as a linear mapping from the initial pressure distribution to the detected acoustic signals.

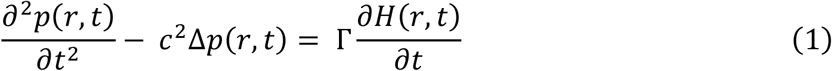

Here, *c* is the speed of sound, *Γ* is the Grueneisen coefficient, and *H* denotes the optical energy deposition per unit volume at position *r* and time *t*.

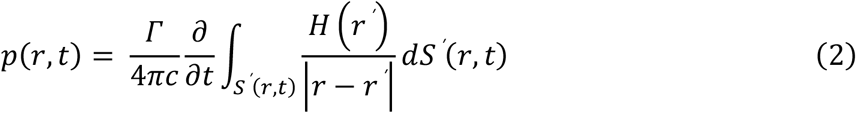

In Eq. (2), the integration is carried out over a time-dependent spherical surface, and the initial condition is defined by the initial pressure distribution generated by the short laser pulse.

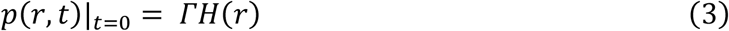

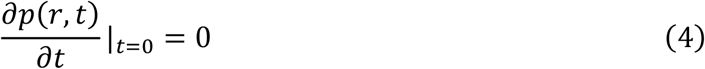

Because the laser pulse duration is much shorter than the acoustic transit time, the source term can be treated as time independent.

To reconstruct the initial pressure distribution in Eq. (3), conventional back-projection methods such as 3D-DAS are computationally efficient but suffer from approximation errors and limited- view artifacts [13, 31-36]. In contrast, 3D model-based reconstruction explicitly accounts for acoustic wave propagation and formulates reconstruction as an inverse problem.

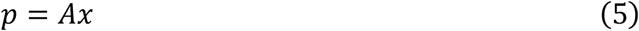

In Eq. (5), *p* denotes the measured photoacoustic signals, *x* represents the spatial distribution of optical energy deposition in vector form, and *A* is the system matrix determined by the imaging geometry and acoustic properties of the medium.

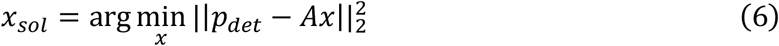

In practice, the inverse problem is solved iteratively using least-squares QR-based iterative method (LSQR) [37]. Because the minimization in Eq. (6) is ill posed under limited-view conditions, particularly in 3D imaging [11, 15-17], regularization is required to stabilize the solution and suppress artifacts.

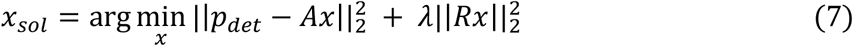

Here, *R* is the regularization matrix, *λ* > 0 is the regularization parameter, and ‖·‖_2_ is the ℓ_2_ norm. The regularization parameter can be selected, for example, using the *L*-curve criterion [38].

### B. Weighted Regional Laplacian Regularization

Incorporating prior structural information into the regularization term can substantially improve reconstruction quality. Conventional Tikhonov regularization promotes global smoothness but does not account for anatomical heterogeneity. To address this limitation, spatially varying regularization strategies have been developed [39]. Instead of using an identity matrix as in conventional Tikhonov regularization, the regularization matrix *R* is defined as a diagonal matrix with voxel-wise weight ±_*i*_, determined by *c*_*i,m*_ and *w*_*m*_, where *c*_*i,m*_ denotes the occupancy fraction of voxel *i* belonging to segment *m*, and *w*_*m*_ represents the regularization strength assigned to segment *m*.

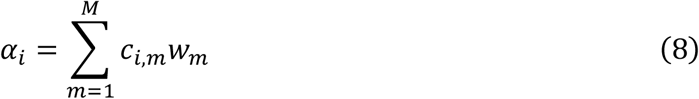

Here, *M* denotes the total number of anatomical segments.

In our work, high-resolution anatomical priors are derived from ULM and incorporated into reconstruction using a weighted formulation. Because ULM has substantially finer resolution than PAT, each PAT voxel is modeled as a mixture of anatomical components defined on the ULM grid through a segmentation matrix.

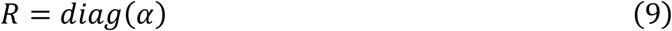

These voxel-wise weights form a vector that is assembled into a diagonal regularization matrix. In this formulation, higher weights impose stronger regularization and suppress variability, whereas lower weights preserve flexibility in regions that are expected to contain fine vascular structure [40].

An alternative strategy, regional Laplacian regularization, promotes intra-segment smoothness while allowing discontinuities across segment boundaries [41].

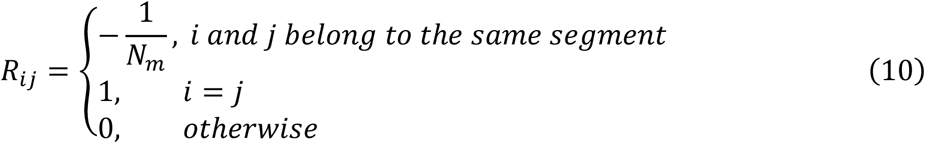

Here, *N*_*m*_ is the number of voxels in segment *m*, assuming exclusive segment membership. This formulation encourages the solution within each segment to approach its segment-wise average. To combine anatomical weighting with intra-segment smoothing, we adopt a weighted regional Laplacian regularization.

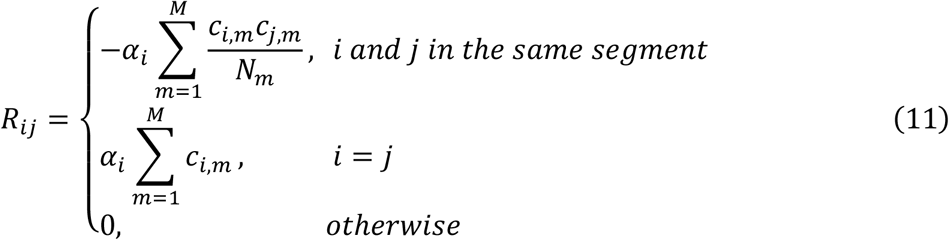

This design applies spatially-varying constraints that promote smoothness within anatomical segments while preserving contrast between different regions. In our implementation, the segmentation consists of two classes—vasculature and background—derived from ULM. This simple representation is sufficient to capture the dominant structural priors for vascular imaging.

### C. Reconstruction Procedure

The reconstruction pipeline of 3D-PAULM^prior^ consists of four stages, as shown in **Fig. 1a**.

**Fig. 1.**
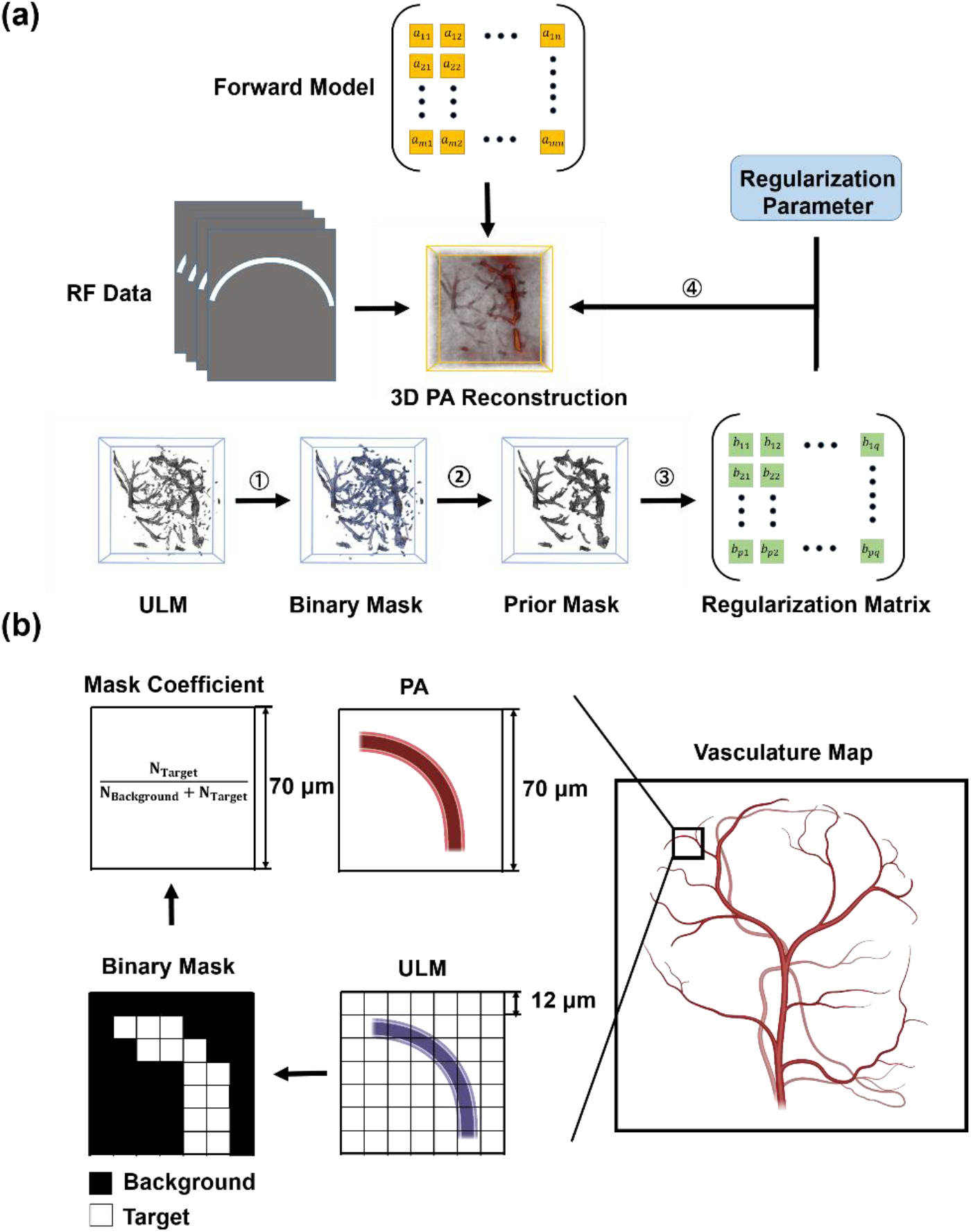
Model-based PA reconstruction framework with ULM prior. (a) Overview of the proposed 3D-PAULM^prior^ reconstruction pipeline. Raw PA signals are reconstructed using a model-based framework regularized by structural priors derived from ULM. The ULM density map is normalized, log-compressed, thresholded, spatially smoothed, and refined by connected- component analysis to generate a vascular prior. This prior is incorporated into a weighted regional Laplacian regularization matrix and used in LSQR-based iterative reconstruction. (b) Construction of voxel-wise mask coefficients. The high-resolution ULM vascular mask is downsampled to the PA reconstruction grid by calculating the vessel occupancy ratio within each PA voxel, yielding a soft vascular prior. Vessel voxels are assigned lower regularization weights to preserve structural detail, whereas background voxels are assigned higher weights to suppress noise.

Step 1. The ULM density map was normalized, log-compressed, and thresholded to generate an initial binary vascular mask. A threshold of −25 dB was selected empirically to preserve vascular structures while suppressing background noise.

Step 2. The binary mask was smoothed using a 3D Gaussian filter (σ = 0.15; kernel size = 3×3×3) to improve spatial coherence, followed by 26-connectivity connected-component analysis to isolate continuous vascular structures.

Step 3. The refined ULM mask was downsampled to the PAT grid by computing the vessel occupancy ratio within each PA voxel, yielding a soft prior. Based on this prior, spatially-varying regularization weights were assigned: lower weights for vascular voxels and higher weights for background voxels. Off-diagonal terms were introduced to enforce intra-region smoothness, as depicted in **Fig. 1b**.

Step 4. The inverse problem was solved using LSQR with a maximum of 100 iterations. The regularization parameter *λ* was selected via *L*-curve analysis to balance data fidelity and prior enforcement [42].

### D. Numerical Simulations

The proposed method was first evaluated using numerical simulations implemented in the k-Wave toolbox [43] and compared with 3D-DAS and Tikhonov-regularized model-based reconstruction (3D-MBT). The simulation domain consisted of 626 × 720 × 720 voxels with an isotropic voxel size of 80 µm, and a perfectly matched layer of 24 grid points was used to suppress boundary reflections. The detector geometry matched the experimental setup with 256 transducer elements.

The imaging target was constructed from an MRA-derived human brain vascular dataset [44]. After segmentation and smoothing, two representative vessels were positioned within a 9.6 × 9.6 × 9.6 mm^3^ region of interest to mimic an artery and a vein. Hemoglobin concentrations were assigned to achieve sO_2_ values of 70% and 96%, respectively. Multispectral simulations were performed at 730, 800, and 870 nm using wavelength-dependent absorption coefficients [45]. Spectral unmixing was applied to estimate oxy- and deoxyhemoglobin concentrations [46], without optical fluence compensation. The ULM prior was simulated using the corresponding binary vascular mask. All simulations and reconstructions were performed on a Windows 11 Pro workstation equipped with an AMD64 processor (3.7 GHz), 200 GB RAM, and an NVIDIA RTX A6000 GPU (48 GB).

### E. Experimental Data Acquisition

All experiments were performed using a custom-designed 2D hemispherical ultrasound transducer array (Imasonics, France) comprising 256 elements with a center frequency of 4 MHz and a −6 dB bandwidth of 45% in both transmit and receive modes, as shown in **Fig. 2a**.

The array provided an 8 × 8 × 8 mm^3^ field of view and was integrated with a programmable ultrasound scanner (Vantage 256, Verasonics, Kirkland, WA) and a Q-switched laser with optical parametric oscillator (Innolas SpitLight I200 OPO, Amplitude Laser) for multispectral PA excitation. During imaging, the transducer array faced upward inside a temperature-controlled water bath, while the imaging target was mounted on an optically and acoustically transparent holder and raster scanned using a 2D motorized stage. To accumulate sufficient frames for ULM and minimize spatial misalignment between PA and US data, 10 cycles of alternating US and PA acquisitions were performed. In the whole-brain imaging, an extended field of view of 12 × 12 × 4.5 mm^3^ was achieved by montaging four scanning positions with a 4 mm step size.

**Fig. 2.**
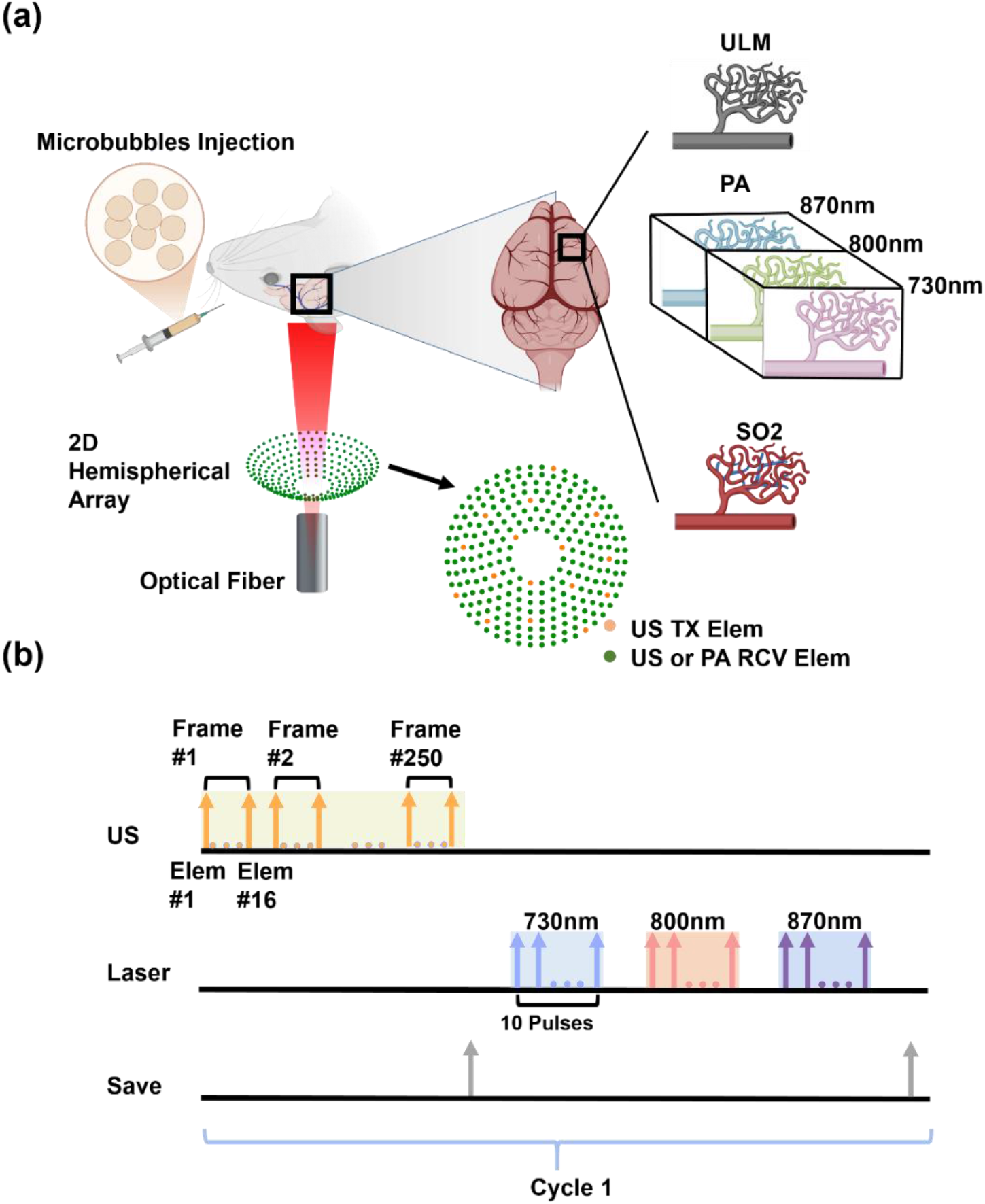
Integrated 3D-PAULM imaging system. (a) Schematic of the hemispherical-array 3D- PAULM imaging system. Synthetic-aperture ultrasound acquisition is performed using 16 transmit events for ULM, while multispectral PA signals are acquired through optical excitation delivered via the central opening of the array. (b) Interleaved PA and US acquisition sequence. PA volumes are acquired at 30 Hz, and US frames are acquired at 403 Hz. Each acquisition cycle includes 250 US frames and 10 PA frames per wavelength, repeated over 10 cycles.

US imaging employed synthetic aperture acquisition with 16 single-cycle transmissions at the transducer center frequency (4 MHz), yielding a frame rate of 403 Hz. Backscattered echoes were sampled at 15.625 MHz by all 256 elements, and 2,500 US frames were acquired at each position. To reduce motion artifacts caused by respiration, a correlation-based rejection method was applied to the radio-frequency data using the averaged signal for each wavelength as the reference; frames with a correlation coefficient below 0.99 were discarded. For ULM reconstruction, B-mode volumes were generated using 3D-DAS with a voxel size of 60 µm and then upsampled by a factor of 5 to a final pixel size of 12 µm, enabling sub-diffraction localization accuracy [47-49].

PA imaging was performed at 730, 800, and 870 nm, scanned sequentially with 10 pulses acquired at each wavelength. Approximately 10% of the laser energy was redirected by a beam splitter to a power meter to record pulse-to-pulse energy fluctuations for optical fluence calibration. Illumination was delivered through an optical fiber bundle (Dolan Jenner; 50% coupling efficiency) inserted through the central opening of the transducer array. All fluence levels remained within ANSI safety limits [50]. PA signals were sampled at 20.83 MHz and averaged across 10 cycles to improve signal-to-noise ratio, as illustrated in **Fig. 2b**.

### F. Phantom Experiment

A pair of closely spaced silicone tubes (inner diameter: 300 µm) was embedded in an agar– intralipid tissue-mimicking phantom (1 mL intralipid per 35 mL solution). One tube was filled with deoxygenated bovine blood prepared by adding sodium hydrosulfite (final concentration, 2.5 mg/mL), whereas the other was filled with oxygenated blood prepared using sodium bicarbonate (final concentration, 37 mg/mL). The tubes were aligned in parallel and connected to a syringe pump to maintain continuous flow and prevent clot formation.

### G. *In vivo* Mouse Brain Imaging

A male mouse with shaved hair was imaged to evaluate 3D-PAULM^prior^ reconstruction. Throughout the imaging session, a catheter was maintained in the femoral vein for microbubble delivery, and body temperature was held at 37 °C. Microbubbles (7 × 10^8^ units/mL) were continuously infused using a syringe pump at 10 µL/min, with a total injection volume below 100 µL. The PA–US dual-mode acquisition lasted approximately 5 min. All animal procedures were approved by the Institutional Animal Care and Use Committee at Duke University.

### H. Image Quality Evaluation

#### 1) Full Width at Half Maximum (FWHM)

Spatial resolution was quantified using full width at half maximum (FWHM), defined as the width of the intensity profile at half of its maximum value [51].

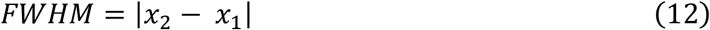

where *P*(*x*_1_) = *P*(*x*_2_) = 0.5 *P*_*peak*_.

#### 2) Contrast to Noise Ratio (CNR)

Contrast-to-noise ratio (CNR) was used to quantify the separability of the region of interest from the background [52].

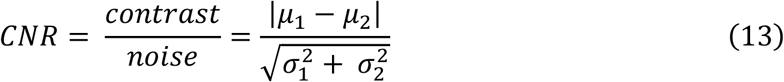

where *μ*_1_, *μ*_2_ and *σ*_1_, *σ*_2_ are the means and standard deviations in ROI and background, respectively.

#### 3) Structural Similarity Index (SSIM)

Structural similarity index (SSIM) was used to quantify structural fidelity relative to the ground truth [53].

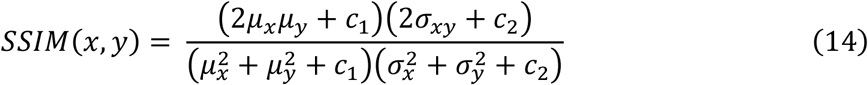

here *μ*_*x*_ and *μ*_*y*_ are the mean intensities of the reconstructed image *x* and the reference image *y* (ground truth), respectively. Similarly, 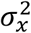 and 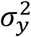 represent their corresponding variances, and 2*σ*_*xy*_ denotes the covariance between *x* and *y*. Constants *c*_1_ and *c*_2_ are used to stabilize the division when the denominator is small.

## Results

We first evaluated the performance of 3D-DAS, 3D-MBT, and 3D-PAULM^prior^ using simulated vascular structures with known oxygenation levels (**Fig. 3a**). Arteries and veins were assigned sO_2_ values of 96% and 70%, respectively. Maximum intensity projections at 800 nm (**Fig. 3b**) show that 3D-PAULM^prior^ most accurately reconstructs vascular morphology and suppresses limited-view artifacts. In contrast, 3D-DAS exhibits severe blurring and structural distortion, whereas 3D-MBT partially improves image quality but still fails to recover fine structures.

**Fig. 3.**
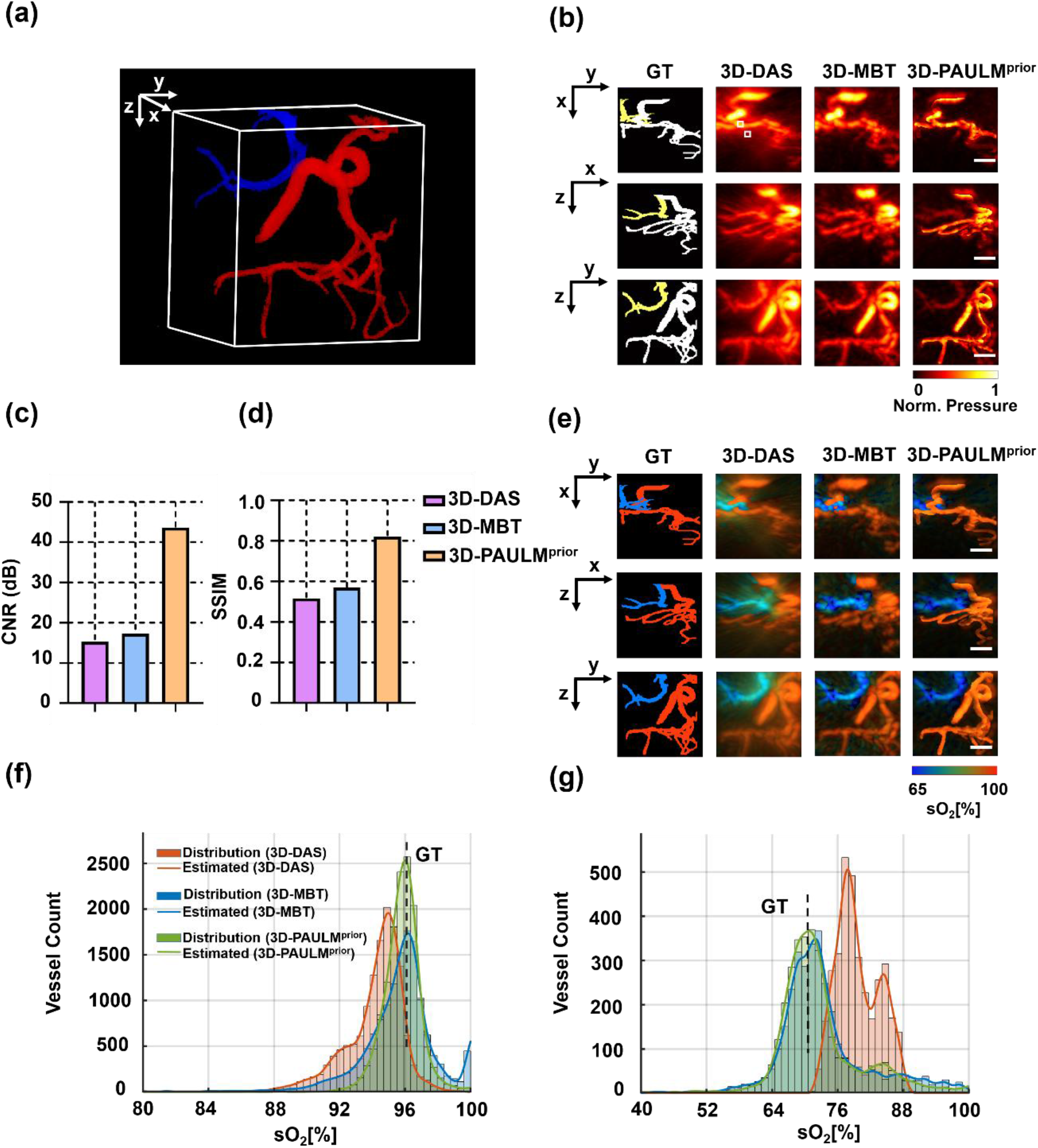
Numerical validation using simulated brain vasculature. (a) Ground-truth vascular model derived from human brain MRA data, with arteries assigned sO_2_ ≈ 96% and veins assigned sO_2_ ≈ 70%. (b) Maximum intensity projections of the ground truth and reconstructed PA images at 800 nm using 3D-DAS, 3D-MBT, and 3D-PAULM^prior^. (c,d) Quantitative comparison of CNR and SSIM. GT, ground truth. (e) Reconstructed sO_2_ maps from multispectral data. (f,g) Histograms and fitted distributions of estimated sO_2_ values in arterial and venous regions. Scale bar: 2.5 mm.

Quantitative evaluation confirms these observations: 3D-PAULM^prior^ achieves the highest CNR (43.67 dB) and SSIM (0.82), substantially outperforming 3D-MBT (17.28 dB, 0.57) and 3D- DAS (15.28 dB, 0.52) (**Fig. 3c, d**).

The advantage of 3D-PAULM^prior^ is further reflected in sO_2_ estimation (**Fig. 3e–g**). The reconstructed distributions closely match the ground truth, with well-separated arterial and venous populations. Specifically, 3D-PAULM^prior^ yields mean sO_2_ values of 95.8% (SD, 1.2%) in arteries and 72.9% (SD, 9.5%) in veins. In contrast, 3D-DAS markedly overestimates venous oxygenation (79.7%) and shows a bimodal venous distribution, indicating instability in spectral unmixing. 3D-MBT improves accuracy but retains increased variance, particularly in venous regions.

We next validated the method using a tissue-mimicking phantom containing two closely spaced tubes filled with oxygenated and deoxygenated blood. As shown in **Fig. 4a, b**, 3D-PAULM^prior^ clearly resolves the two tubes across the imaging depth, whereas 3D-DAS fails to separate them and 3D-MBT shows depth-dependent degradation. These differences become more pronounced away from the acoustic focus, where conventional reconstructions are increasingly affected by artifacts.

**Fig. 4.**
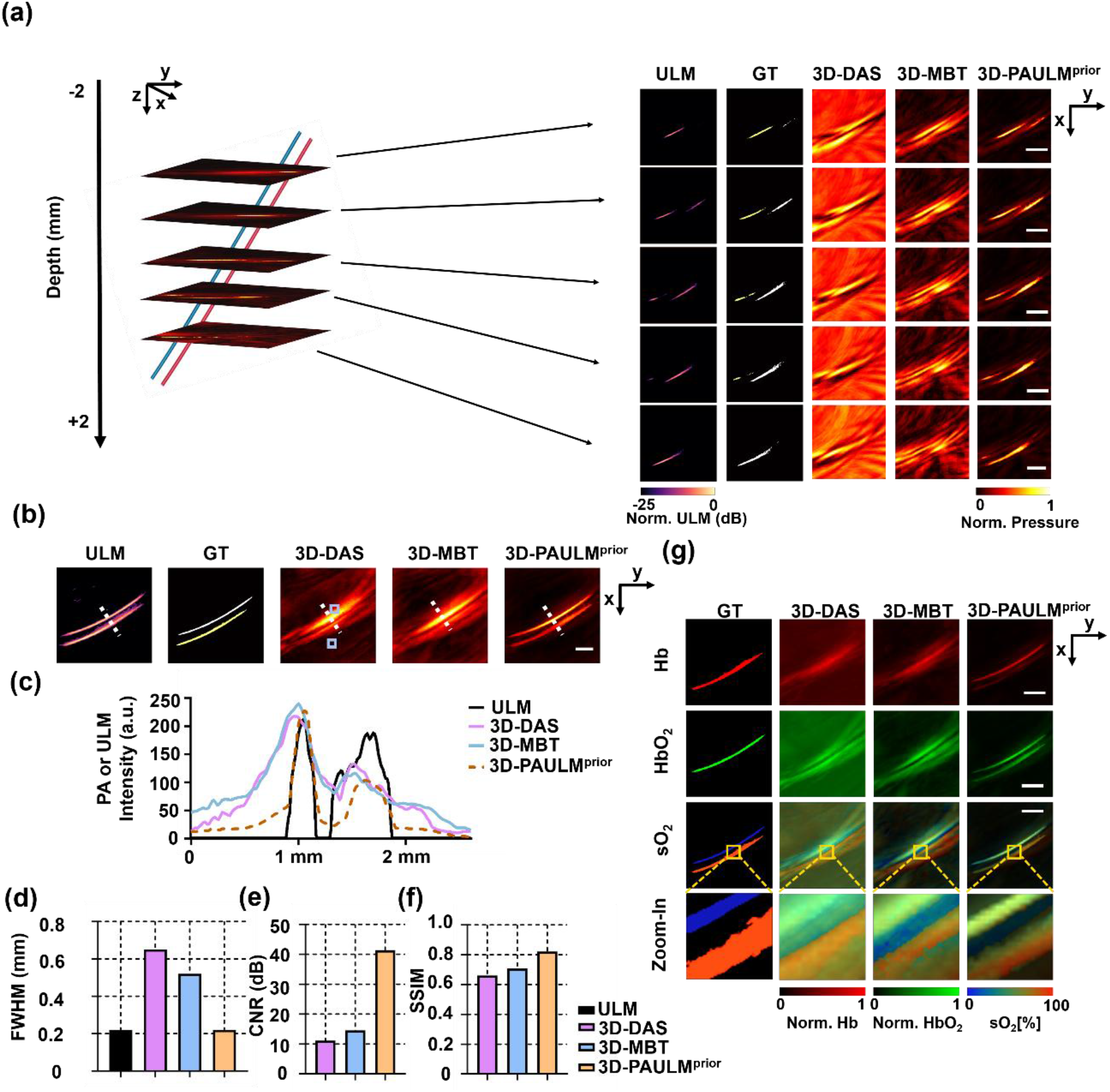
Phantom validation using paired oxygenated and deoxygenated blood tubes. (a) Cross-sectional ULM and PA reconstructions at 800 nm over depths from −2 mm to 2 mm, with 0 mm corresponding to the center of the field of view. (b) *x–y* maximum intensity projections of the ULM, ground truth, and reconstructed PA images. (c) Line profiles extracted along the dashed lines in (b). (d–f) Quantitative comparison of FWHM, CNR, and SSIM. (g) Reconstructed HbR, HbO_2_, and sO_2_ distributions from different reconstruction methods. Scale bar: 2.5 mm.

Line-profile analysis (**Fig. 4c**) and FWHM measurements (**Fig. 4d**) show that 3D-PAULM^prior^ achieves a spatial resolution of 0.22 mm, improved spatial resolution by over 50% compared with 3D-DAS. Quantitatively, 3D-PAULM^prior^ improves CNR by 261.2% and SSIM by 24.6% relative to 3D-DAS (**Fig. 4e,f**). Functional imaging results (**Fig. 4g**) further highlight the advantage of 3D-PAULM^prior^: the reconstructed sO_2_ map shows a sharp transition between the two tubes and closely matches the expected values. In contrast, 3D-MBT produces discontinuities, whereas 3D-DAS shows substantial mixing between the two regions.

We further evaluated 3D-PAULM^prior^ *in vivo* using mouse brain imaging (**Figs. 5 and 6**). Compared with 3D-DAS and 3D-MBT, 3D-PAULM^prior^ provides substantially improved visualization of cerebrovascular structures. As shown in **Fig. 5a**, 3D-DAS yields diffuse and poorly resolved images, while 3D-MBT partially recovers vascular structures. In particular, vertically oriented vessels are largely absent in both methods owing to limited-view constraints. By incorporating ULM priors, 3D-PAULM^prior^ effectively recovers these missing structures and produces a more complete vascular network.

**Fig. 5.**
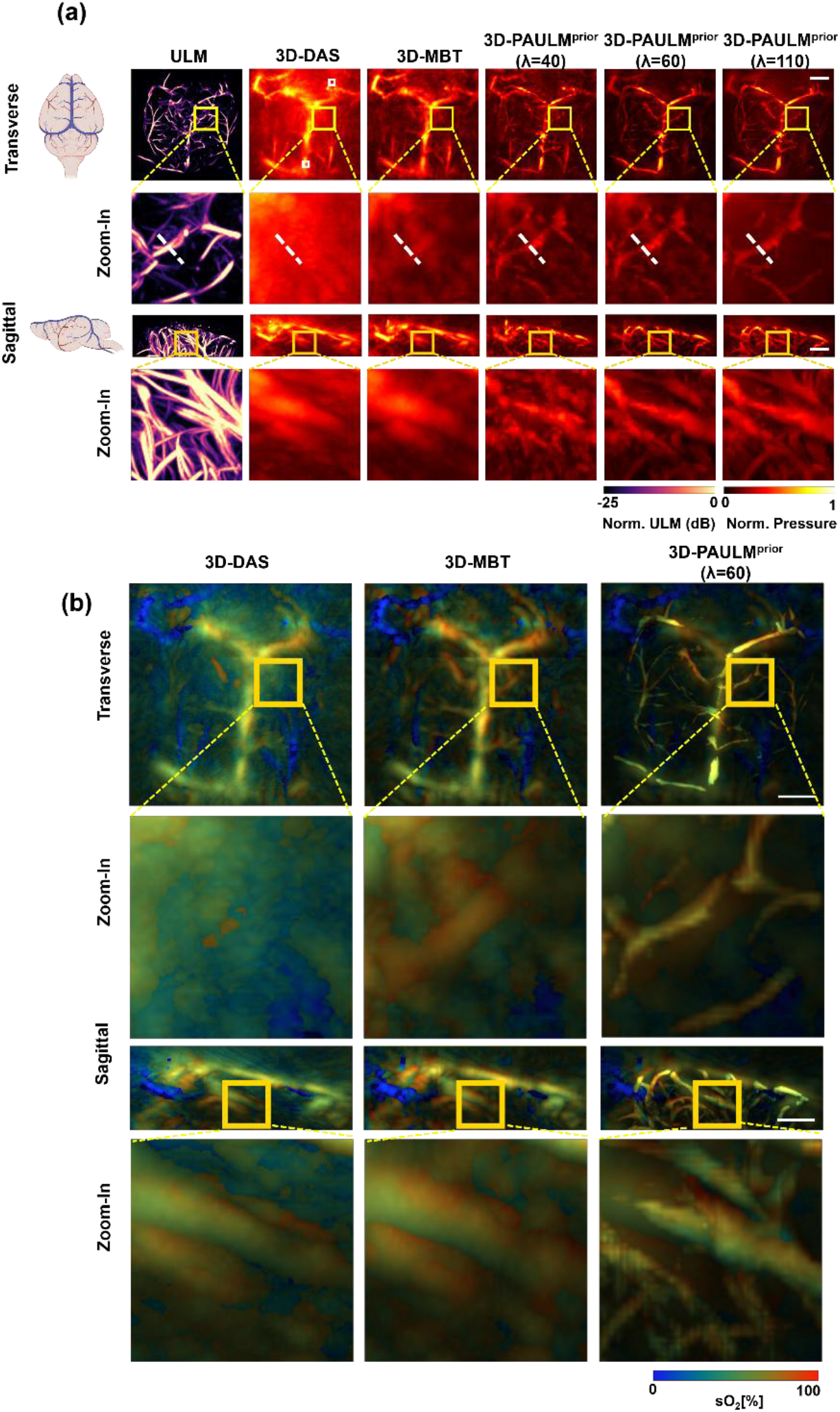
*In vivo* mouse brain imaging. (a) Transverse and sagittal maximum intensity projections of ULM and PA reconstructions using 3D-DAS, 3D-MBT, and 3D-PAULM^prior^ with different regularization strengths. (b) Reconstructed sO_2_ maps from multispectral PA imaging using 3D- DAS, 3D-MBT, and optimized 3D-PAULM^piror^.Scale bar: 2.5 mm.

**Fig. 6.**
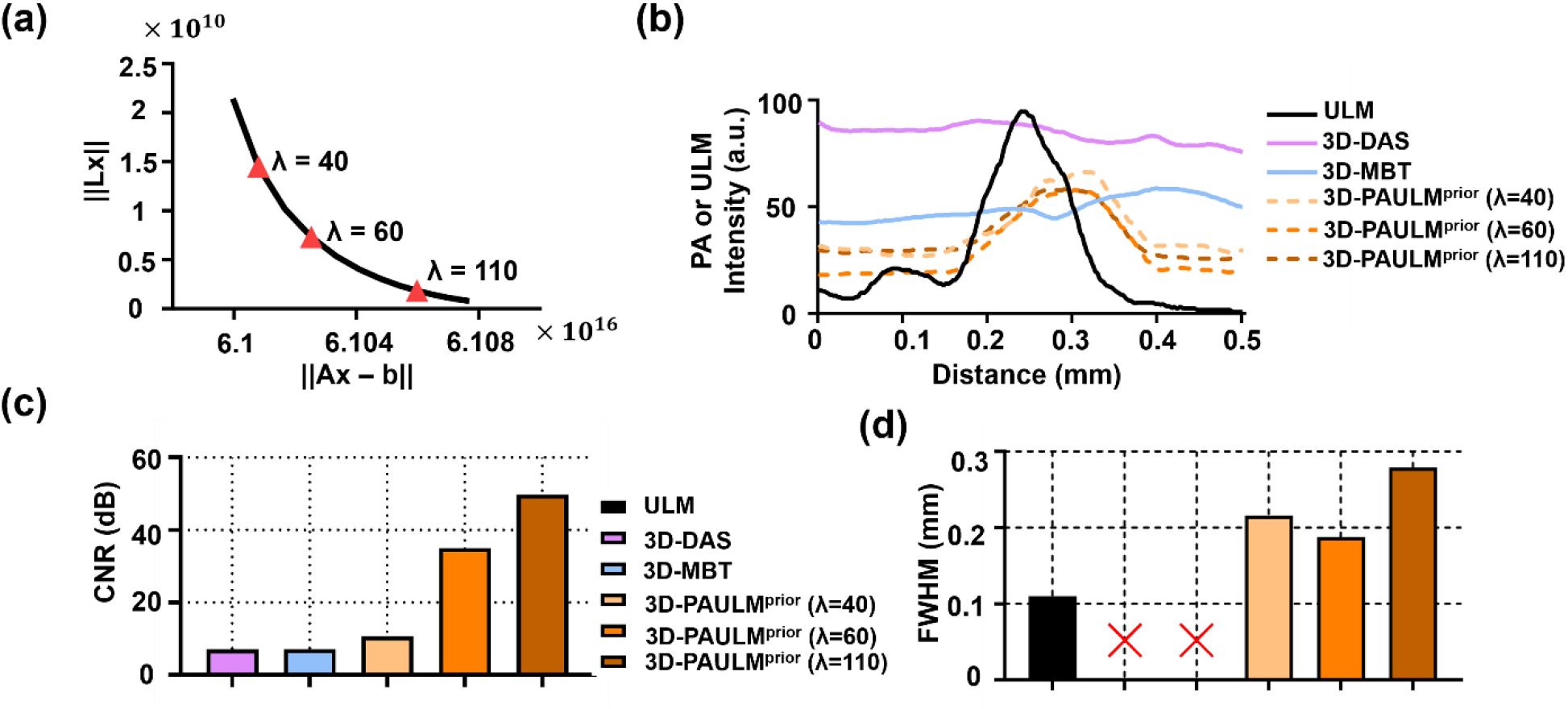
Quantitative comparison of *in vivo* reconstruction performance. (a) *L*-curve analysis for selecting the regularization strength in 3D-PAULM^prior^. (b) Line profiles extracted from the zoomed-in transverse views in Fig. 5a. (c) CNR measured within the ROIs marked in Fig. 5a. (d) FWHM derived from the line profiles in (b). FWHM values for 3D-DAS and 3D-MBT are not reported because they do not resolve the selected vessel.

We also investigated the effect of regularization strength. Increasing λ enhances vessel– background separation and suppresses noise, but excessive regularization leads to over-smoothing and loss of fine structures. Among the tested values, λ = 60 achieves the best balance between resolution and contrast, as confirmed by the *L*-curve analysis (**Fig. 6a**). Quantitatively, 3D-PAULM^prior^ (λ = 60) achieves the highest CNR and the narrowest FWHM among all PA reconstructions (**Fig. 6c, d**). Multispectral sO_2_ maps (**Fig. 5b**) further demonstrate improved spatial confinement and reduced background contamination relative to 3D-DAS and 3D-MBT. The resulting oxygenation maps show clear vessel-specific contrast and physiologically consistent gradients.

## Discussion and Conclusion

In this work, we introduced 3D-PAULM^prior^, a model-based reconstruction framework that integrates super-resolved vascular priors from ultrasound localization microscopy into photoacoustic imaging. By combining weighted regional Laplacian regularization with co-registered ULM data, the method addresses key limitations of conventional PAT reconstruction, including diffraction-limited resolution, limited-view artifacts, and instability in functional quantification.

Across simulations, phantom experiments, and *in vivo* imaging, 3D-PAULM^prior^ consistently improves both structural and functional performance. The method enhances vessel delineation, suppresses background artifacts, and redistributes energy toward true absorbers, resulting in higher CNR, improved SSIM, and better resolution. Notably, the achieved resolution approaches that of ULM in controlled settings while preserving the penetration depth and functional sensitivity of PAT.

A central advantage of 3D-PAULM^prior^ is its ability to reduce limited-view artifacts. By incorporating ULM-derived structural priors, the method compensates for missing angular information and recovers vascular features that are otherwise poorly reconstructed. Importantly, the ULM prior is used as a soft regularization constraint rather than a hard anatomical mask; the reconstructed optical properties therefore remain governed by the measured PA signals, while the prior guides the solution toward anatomically plausible vascular structures. This distinction matters because ULM and PA encode related but non-identical vascular information: ULM reflects microbubble-accessible perfused vessels, whereas PA reflects hemoglobin absorption. Nevertheless, the integration of anatomical priors improves structural fidelity and constrains spectral unmixing to vascular regions, leading to more spatially specific and physiologically consistent sO_2_ maps.

Several limitations remain. First, the current forward model does not account for the transducer electrical impulse response or spatial impulse response, both of which can introduce distortion in wideband or deep-tissue imaging [54-57]. Incorporating these effects should further improve reconstruction accuracy. Second, acoustic heterogeneity is not explicitly modeled. Variations in speed of sound can introduce phase errors and degrade image quality [58, 59]; integrating spatially varying acoustic parameters would improve robustness. Third, the present *in vivo* study serves primarily as a technical proof of concept rather than a statistically powered biological study, and broader validation in larger cohorts will be needed.

In summary, 3D-PAULM^prior^ provides a robust and scalable framework for high-resolution, quantitatively accurate photoacoustic imaging. By leveraging the complementary strengths of ULM and PAT, the method bridges the gap between vascular resolution and functional imaging depth and offers a promising strategy for multimodal vascular imaging in both preclinical research and future clinical applications.

## Acknowledgments

This work was partially sponsored by the United States National Institutes of Health (NIH) grants R01EB037095, R01ES036951, RF1 NS115581, R01 NS111039, R01 EB028143, R01 DK139109, R01 DK052985, R01 MH135932; The United States National Science Foundation (NSF) CAREER award 2144788; American Heart Association Collaborative Science Award (25CSA1417550); Duke Gilhuly Acceleration Grant; Duke University Pratt Beyond the Horizon Grant; Chan Zuckerberg Initiative Grant (2024-349531); Duke University DST Spark Seed Grant; Duke Coulter Translational Grant; North Carolina Biotechnology Center Triangle Research Grant (2024-TRG-0041).

## Competing interests

J.Y. has a financial interest in Lumius Imaging, Inc., which did not support this work. The other authors declare no competing interests.

## Notes

### Competing Interest Statement

The authors have declared no competing interest.

